# Dorsal anterior cingulate and ventromedial prefrontal cortex have inverse roles in both foraging and economic choice

**DOI:** 10.1101/046276

**Authors:** Amitai Shenhav, Mark A. Straccia, Matthew M. Botvinick, Jonathan D. Cohen

## Abstract

Recent research has highlighted a distinction between sequential foraging choices and traditional economic choices between simultaneously presented options. This was partly motivated by observations in Kolling et al. (2012) [KBMR] that these choice types are subserved by different circuits, with dorsal anterior cingulate (dACC) preferentially involved in foraging and ventromedial prefrontal cortex (vmPFC) preferentially involved in economic choice. To support this account, KBMR used fMRI to scan human subjects making either a foraging choice (between exploiting a current offer or swapping for potentially better rewards) or an economic choice (between two reward-probability pairs). This study found that dACC better tracked values pertaining to foraging, while vmPFC better tracked values pertaining to economic choice. We recently showed that dACC’s role in these foraging choices is better described by the difficulty of choosing than by foraging value, when correcting for choice biases and testing a sufficiently broad set of foraging values (Shenhav et al., 2014). Here, we extend these findings in three ways. First, we replicate our original finding with a larger sample and a task modified to address remaining methodological gaps between our previous experiments and that of KBMR. Second, we show that dACC activity is best accounted for by choice difficulty alone (rather than in combination with foraging value) during both foraging and economic choices. Third, we show that patterns of vmPFC activity, inverted relative to dACC, also suggest a common function across both choice types. Overall, we conclude that both regions are similarly engaged by foraging-like and economic choice.

## Introduction

Foraging is a type of decision-making characterized by a series of sequential choices between a default/exploitative option (stay in current patch) and the option of switching to another patch. Research has grown increasingly interested in the computational and neural basis for modern-day analogs of foraging decisions (Blanchard & Hayden, 2015; Calhoun & Hayden, 2015; Carter, Pedersen, & Mccullough, 2015; Constantino & Daw, 2015; Hayden, Pearson, & Platt, 2011; Hayden & Walton, 2014; Kolling, Behrens, Mars, & Rushworth, 2012; Kolling, Behrens, Wittmann, & Rushworth, 2016; Kolling, Wittmann, & Rushworth, 2014; Kvitsiani et al., 2013; Rushworth, Kolling, Sallet, & Mars, 2012). In particular, recent studies have contrasted foraging decisions with traditional *economic* decisions, which are typically characterized by choices between multiple simultaneously presented options that offer different tradeoffs between risk and reward. Recently, Kolling and colleagues (Kolling et al., 2012; Kolling et al., 2016; Kolling & Hunt, 2015; Rushworth et al., 2012) proposed that this may represent a fundamental distinction that is realized neurally, with different brain regions having been evolved to drive decision-making in these two choice settings (see also Boorman, Rushworth, & Behrens, 2013; Kolling et al., 2014). Specifically, these authors suggest that dorsal anterior cingulate cortex (dACC) and ventromedial prefrontal cortex (vmPFC) may have evolved to play critical and differential roles in foraging versus economic choice, respectively.

To support this proposed dichotomy, Kolling and colleagues (2012 [hereafter, KBMR]) designed a task that alternates between offering participants opportunities to make foraging choices (stick with a set of rewards offered to you or swap for potentially better rewards, at the cost of time and lost reward) and economic choices (choose between two immediately available probabilistic rewards). This study reported that dACC tracked the relative value of foraging (RVF) during the foraging stage of a trial (Stage 1). Furthermore, they argued that this effect could not be accounted for by the difficulty of making a choice (with which this region has previously been associated; Blair et al., 2006; Pochon, Riis, Sanfey, Nystrom, & Cohen, 2008; Shenhav & Buckner, 2014), but rather that choice difficulty effects could reflect valuation of a foraging alternative (e.g. the non-default or unchosen option). We recently reported two experiments, showing in the first that dACC activity in KBMR's task could in fact be accounted for by choice difficulty when the measure of difficulty corrected for a participant's default bias against foraging (Experiment 1) and, in the second, that testing a wide enough range of RVFs to dissociate difficulty from RVF in fact reveals an effect of difficulty but not RVF in dACC (Experiment 2) (Shenhav, Straccia, Cohen, & Botvinick, 2014). We interpreted these results as suggesting that dACC's role in foraging was representative of a more general and consistent role in valuation and cognitive control, rather than this region instead being primarily concerned with foraging valuation.

Here we extend these findings in two ways. First, we perform a stronger test of an alternative model of dACC whereby activity in this task reflects *both* choice difficulty and RVF rather than one or the other. We do this by testing a larger sample of participants and by closing gaps between the designs of our earlier studies and that of KBMR that may have reduced our sensitivity to a foraging value signal that rode atop a difficulty signal (cf. Kolling et al., 2016). In particular, we now employ reward-related stimuli that are multidimensional, and we enforce a delay before participants are able to respond. Second, we explore whether the role of vmPFC in foraging versus economic choice is fundamentally different (as proposed in KBMR) or whether, like dACC, a single variable can capture the vmPFC’s role in both choice settings. Specifically, given the modifications just described, we test whether vmPFC tracks relative chosen value for both foraging and economic choices.

We replicate our previous findings in dACC, showing that this region continues to only track choice difficulty rather than both difficulty and foraging value, even with modifications to our task and a larger sample. Furthermore, using our modified task design, we find that vmPFC’s role in the task is indeed similar across both stages of the task, and inverse to that of the dACC; vmPFC BOLD activity tracks the relative ease of choosing (or the magnitude of the difference between one’s options).

## Method

### Participants

We recruited 34 neurologically healthy, right-handed participants (30 female; M_age_ = 21.3, SD_age_ = 2.9) to participate in a neuroimaging experiment. Of these, three participants were excluded: one for misunderstanding task instructions (assuming that Stage 2 could be predicted from success at Stage 1), one for never foraging in the first half of the session (the participant reported switching evaluation strategies at that point), and one for excessive head movement. This yielded datasets from 31 participants (27 female; M_age_ = 21.3, SD_age_ = 2.8) for our final analysis. For three of those datasets, one of the three task blocks was excluded, in one case for excessive head movement and in the other two cases for falling asleep during that block.

### Foraging task

Participants performed a task based on one introduced in KBMR (and used in modified form by Shenhav et al., 2014) to simulate foraging-like choice (Figure 1A). Briefly, each trial is made up of two stages. In Stage 1 (foraging stage), participants are offered two potential rewards (the engage option), shown in the middle of the screen, which can be carried with them directly to Stage 2 (the engage stage). If they choose to keep this engage option and move to Stage 2, they are then offered a probabilistic choice between those two potential rewards (with probabilities only revealed at the start of Stage 2); are paid based on the outcome of the gamble they select; and then move on to Stage 1 of a new trial. Before choosing to move to Stage 2, however, participants can choose to swap the two rewards in their current engage option with two rewards chosen at random from a set of six potential alternatives shown in a box at the top of the screen (the forage set). They can choose to do this as many times as they prefer during Stage 1, but each time they do so they incur a search cost (indicated by the color of the box) and wait a variable delay period (see below) before the new engage and forage options appear. Rewards received at the end of Stage 2 accumulate across trials towards a fixed target appearing at the bottom of the screen (reward bar not shown in Figure 1), and participants received a monetary payoff at the end of the session proportional to the number of times the reward bar reached this target and reset.

In Shenhav et al. (2014), we performed two experiments that maintained this task structure; one included a relatively narrow range of forage values across trials (largely favoring decisions to engage; Experiment 1), replicating KBMR; the other included a much wider range of forage values (Experiment 2). In order to collect response times, both experiments allowed participants to respond as soon as the relevant choice variables appeared on the screen for a given stage, a deviation from KBMR’s original study wherein participants waited a variable delay before they were allowed to respond in each trial stage. Experiment 2 further deviated from KBMR’s task by utilizing numeric stimuli to represent the (now much wider) range of rewards, rather than using a limited set of symbols with learned values (as in Experiment 1).

The task in the current study was identical to the one used in Experiment 2 of this earlier study but avoided either of the deviations just described. First, rather than allowing participants to respond immediately upon viewing their options, the same variable time delay was imposed during both choice stages as in KBMR. Specifically, once the options appeared participants waited an average of 2.7s and 1.8s in Stages 1 and 2, respectively, before a question mark appeared in the center of the screen informing them that they can respond when ready (these delays were Poisson-distributed from 2–4s and 1–4s ranges, with additional 200ms jitter added for design efficiency). While this prevented us from recording decisions that occurred within this delay period-resulting in response times that were noisier and potentially overestimated the true decision time, relative to a traditional free-response paradigm-it ensures that participants are
presented with the options for a minimal period before responding, potentially increasing our sensitivity to neural activity associated with evaluation of those options (particularly for fast choice trials).

Second, in place of purely numeric stimuli that indicate exact point values for each of the items in the foraging and engage set, we followed KBMR’s practice of using more abstract symbols. Specifically, we used playing cards to represent potential reward values ranging from 20 to 530 points. The value of each card was indicated both by its suit and the number or face that appeared on it. Within a suit, reward values increased as follows: 2–10, Jack, Queen, King, Ace (1). Across suits, values increased as follows: clubs, diamonds, hearts, spades. Participants were told these reward mappings and then tested on them extensively, first by performing the same binary choice task used in KBMR to criterion (95% accuracy) and then sorting a random subset of the deck in the appropriate order. We chose playing cards as our stimuli in order to increase integration demands (across stimulus dimensions) relative to entirely numeric stimuli, while still providing the necessary range of values to dissociate our variables of interest without the burden of learning the values of many more novel stimuli than the twelve that were used in the original study. Moreover, as in KBMR and Experiment 1 of Shenhav et al., 2014, search costs were only indicated by the color of the search box, with green/white/red corresponding to point losses of 5/20/40. (Also, as in KBMR, search costs were incurred probabilistically-on 70% of forages-and the outcome of this gamble was revealed after a given choice to forage.) Collectively these factors made it difficult for a participant to engage in arithmetic calculations to determine their choice (particularly given the differential weighting of option values that would have been required to produce the observed choice behavior; see Figure 1B).

As in Experiment 2 of our previous study, we configured the values for the (initial) foraging stage of each trial such that they covered a wide distribution of relative forage values (i.e., balanced over the full range of relative advantages of foraging vs. engaging and vice versa). A total of 200 trials were generated, with relative forage values sampled approximately uniformly from the range –307 to 458 (where these values reflect the difference between the average of the forage and engage sets on a given trial; see below for details of how forage values were calculated for our primary analyses). Note that the final number of foraging choices and distributions of values across them was a combination both of these initial 200 choices as well as the new choices that were generated each time the participant chose to forage; these new choices consisted of the previous choice set with a swap of the engage values and a random subset of two forage values. The number of Stage 2’s (i.e., opportunities to harvest reward) was fixed at 200 across subjects.

The remaining timing details were identical to KBMR's task. The average intertrial intervals (ITIs) was 3.0s between choosing to forage and receiving search cost feedback (range: 2–6s); 2.7s between foraging feedback (1–2s duration) and the next Stage 1 choice set for that trial (range: 2–4.5s); 4.5s between Stage 1 choices to engage and being shown the engage probabilities (range: 3–8s); 4.5s between Stage 2 choices and reward feedback (range: 3–8s); and 2.7s between Stage 2 feedback and Stage 1 of the next trial (range: 2–4.5s)

### Image acquisition

Scanning was performed on a Siemens Skyra 3T MR system. Similarly to KBMR and our previous experiment, we used the following sequence parameters for the main task: field of view (FOV) = 196mm x 196mm, matrix size = 66 x 66, slice thickness = 3.0mm, slice gap = 0.0mm, repetition time (TR) = 3.0s, echo time (TE) = 30ms, flip angle (FA) = 87°, 54 slices, with slice orientation tilted 15° relative to the AC/PC plane. At the start of the imaging session, we collected a high-resolution structural volume (MPRAGE) with the following sequence parameters: FOV = 200mm x 200mm, matrix size = 256 x 256, slice thickness = 0.9mm, slice gap = 0.45mm, TR = 1.9s, TE = 2.13ms, FA = 9°, 192 slices.

### Behavioral analysis

In order to avoid strong assumptions about how participants represented the relative value of foraging, we used a “model-free” approach to inferring these values from the behavioral data.

We performed a logistic regression on a given participant’s choices, against the (normalized) variables that were potentially relevant to their decision (the same procedure used in KBMR for behavioral but not fMRI analyses; shown in Figure 1B). Based on these within-subject logistic regression coefficients and the values presented on a given trial, we generated predictions of the likelihood (log-odds) of choosing one option or the other. We use this log-odds ratio as our primary estimate of the relative value of foraging (RVF) on a given trial, and take its absolute value (i.e., distance from indifference) as an estimate of the ease of choice on that trial (see also Supplementary Figure 5C of Shenhav et al., 2014). We generated analogous estimates of relative value (RV) and difficulty in Stage 2 based on a regression of choices onto the magnitudes and probabilities of reward for the left and right option in a given choice. Because these estimates correct for subject-specific bias and option weighting, we will occasionally also refer to them as RVFc and RVc.

Where relevant, we also report the outcome of analyses using RV estimates comparable to those used by KBMR that are uncorrected in these respects (RVkbmr). In Stage 1 these include an RVF estimate that compares the search set and the engage pair, using a weighting function that prioritizes the highest engage value (see description of ‘search evidence’ in KBMR and our previous paper); a relative chosen value estimate (RV_Chosen-KBMR_) that compares the average of the search set (search value) to the search cost and the weighted average of the engage pair (engage value), with each variable signed according to whether the forage or engage option was chosen. In Stage 2, RV_Chosen-KBMR_ consisted of the difference between the expected values (reward multiplied by probability) of the chosen and unchosen options.

### fMRI analysis

All neuroimaging analyses were based on those reported in KBMR and Shenhav et al. (2014). Imaging data were analyzed in SPM8 (Wellcome Department of Imaging Neuroscience, Institute of Neurology, London, UK). Functional volumes were motion corrected, normalized to a standardized (MNI) template (including resampling to 2mm isotropic voxels), spatially smoothed with a Gaussian kernel (5mm FWHM), and high-pass filtered (0.01 Hz cut-off).

Our subject-level GLMs modeled separate regressors for the decision and feedback phases of Stages 1 and 2. All regressors were modeled as stick functions, and additional nuisance regressors were included for motion within each block. Decision phase regressors were further modulated by non-orthogonalized parametric regressors, as described below.

Two GLMs were aimed at exploring the simple effect of RVF (in the absence of additional covariates) when focusing on different subsets of the data:

GLM-1: Regressors for RVFc in Stage 1 and RVchosen-c in Stage 2 (Figures 2A and 4A–C). This GLM was used in a sliding window analysis aimed at examining foraging value effects within specific subsets of our data (e.g., to test whether a positive effect of foraging value is found in a range of RVFs comparable to those tested in KBMR). Specifically, we rank-ordered all trials by RVFc and ran a series of GLMs that each tested for a parametric effect of RVFc within a given percentile range window (for example, 0th–50th percentiles). For each of these GLMs, a separate unmodulated regressor modeled all foraging decisions outside of that window. We performed two sets of windowed analyses, one using windows of 70% per GLM, the other using windows of 50% per GLM. Consecutive windows were shifted by 10 percentiles, resulting in 86% and 80% overlap between consecutive windows in the respective analyses.

GLM-2: Regressors for RVFc in Stage 1, separately for trials where subjects chose to forage versus engage, and for RV_Chosen-C_ in Stage 2 (Figures 2B and 3A). A second version of this GLM (GLM-2B) used RVFkbmr in place of RVF (Supplementary Fig. 6A).

GLM-3: Regressors for each of eight equally sized RVF bins. This GLM was used to complement GLM-1; whereas GLM-1 measured the slope of the BOLD-RVF relationship using different subsets of trials, this GLM instead uses discrete bins to capture the shape of the RVF response function across all trials and RVF values. We performed a second version of this analysis (GLM-3B) using RVFkbmr in place of RVF (Supplementary Fig. 6B).

A third GLM was constructed in order to replicate analyses previously used to differentiate roles of dACC and vmPFC in the foraging task:

GLM-4: Regressors for RVF_kbmr_ and search costs in Stage 1, and regressors for RV_Chosen-KBMR_ and log RT in both stages (Figures 5A–B). This GLM focused on the 0^th^–70^th^ percentile range of RVFs, in order to approximate the range of trial values in KBMR.

The remaining GLMs were aimed at comparing the roles of dACC/vmPFC in foraging value (RVF_C_) versus difficulty or relative chosen value (|RVF_C_| or RV_Chosen-C_):

GLM-5: Regressors for RVFc and |RVFc| in Stage 1, RVRight-C and |RVRight-C| in Stage 2 (Figures 2C and 3B). Log RT was also included at both stages for this GLM, with the exception of a second version (GLM-5B) used to re-analyze Shenhav et al. (2014) (this study omitted RT in comparable analyses). A third version (GLM-5C) was identical to GLM-5 but RVF-related regressors excluded the easiest choices (|RVF|>4.5). A fourth version (GLM-5D) included all trials but used a version of RVF that excluded subject-specific bias terms (RVFnoBias).

GLM-6: Regressors for RV_Chosen-C_ and log RT in both stages (Figure 5C).

GLM-7: Regressors for RVF_KBMR_, search costs, and |RVF_C_| in Stage 1; regressor for RVChosen-KBMR in Stage 2; and regressors for log RT in both stages.

GLM-8: Regressors for search value, engage value, search costs, and |RVFc| in Stage 1; regressor for RV_Chosen-KBMR_ in Stage 2; and regressors for log RT in both stages.

For ease of comparison with KBMR and Shenhav et al. (2014), all whole-brain *t*-statistic maps are depicted at a voxelwise threshold of *p* < 0.01, cluster extent thresholded to achieve a family-wise error (FWE) corrected *p* < 0.05. To guard against concerns of Type 2 errors, we also include uncorrected equivalents of key maps in the supplementary materials. All maps are projected onto the Caret-inflated cortical surface (Van Essen, 2005).

Region-of-interest (ROI) analyses examined average beta estimates from spheres (9mm diameter) in dACC and vmPFC. The main dACC ROI was taken from Shenhav et al. (2014), drawn around the foraging value peak in its Experiment 1 (MNI coordinates: x=4, y =32, z=42). To provide a strong within-subject test of foraging value effects, we created a second dACC ROI around the RVF_C_ peak in the current study (x=-6, y=18, z=46), during the 10^th^–60^th^ percentile window in the sliding analysis described above (this is the range for which we find the strongest evidence of a positive RVF effect). We created an analogous ROI within vmPFC around the peak inverse foraging value effect within that same window (x=8, y=32, z=-8).

### Follow-up experiment: Math task

In order to test whether participants performing the foraging task were likely to be using explicit mathematical operations to make their choice (i.e., by averaging over card values and subtracting those averages from one another), we ran a follow-up experiment in which an independent group of participants (N=14) were shown a stimulus array analogous to the one shown in the foraging task, but entirely numeric (Supplementary Figure 8A). They were instructed to average six numbers shown at the top of the screen, subtract from that the middle number and the average of the bottom two numbers, and report the result. They were told that they could round the result of each operation to the nearest whole number. Paralleling the foraging task, numbers at the top and bottom of the screen for a given trial were randomly drawn (without replacement) from the numbers 2–53, and the middle number from the numbers 1–3. Participants typed their response on a keyboard and were given feedback as to whether the response was within ±2 of the correct response. There was no response deadline. For analysis purposes, response times were recorded as the time of the first keystroke in a response. Participants performed three practice trials, and then performed the main task for 60 trials or 45 minutes, whichever came first.

Note that this task provides a conservative comparison to the main foraging task in at least two ways. First, the task replaced symbolic stimuli (card suits/ranks for foraging reward stimuli and frame color for foraging cost) with explicit numbers, and scaled the associated values by an order of magnitude (2–53 rather than 20–530), avoiding any additional computation time incurred by these additional operations. Second, the task instructed participants to weigh the averages of the top, middle, and bottom of the screen uniformly. Participants performing the foraging task weighed these quantities differently (Figure 1B), an additional operation that presumably would have increased the cognitive demands of the math task.

## Results

### Behavioral results

As in earlier studies, a logistic regression revealed that Stage 1 (foraging) choices were sensitive to the average value of the foraging set, the search cost, and the value of the two engage options (Figure 1B; all p<0.001). In addition, subjects exhibited a significant default bias towards engaging rather than foraging (M = –1.4, SE = 0.26, *t*(30) = –5.4, *p*<0.001).

We quantify the relative value of foraging (RVF) value based on the predicted log-odds of foraging versus engaging, as estimated by the within-subject logistic regression above, accounting for subject-specific weights on each of the foraging variables as well as any implicit bias towards engaging (Figure 1C; Supplementary Figure 1A). We use the absolute value of this variable (|RVF|) as our estimate of choice ease/difficulty, with the most difficult trials being closest to indifference and the easiest trials having a high log-odds of either foraging or engaging (see, e.g., Bonnelle, Manohar, Behrens, & Husain, 2015). The distribution of forage values across trials produced a sufficient number of choices with high relative foraging values (i.e., easy choices in favor of foraging) so as to avoid a confound between foraging value and difficulty (average correlation between RVF and |RVF|: Pearson’s *r* = 0.032, average R^2^ = 0.054, sign-rank *p* = 0.91; average Spearman’s *ρ* = 0.17, sign-rank *p*<0.001).

Additional analyses confirmed that our use of playing cards as stimuli did not result in significantly different behavioral patterns than previous studies that used KBMR’s original stimuli (Supplementary Figure 7). Furthermore, this behavior was substantially different than that of participants in a follow-up experiment who were instructed to use explicit mathematical operations to determine foraging value (e.g., decisions on this math task took over twenty times longer to make than foraging choices; Supplementary Figure 8), suggesting that participants were unlikely engaging in such operations to make their foraging choices.

**Figure 1.**
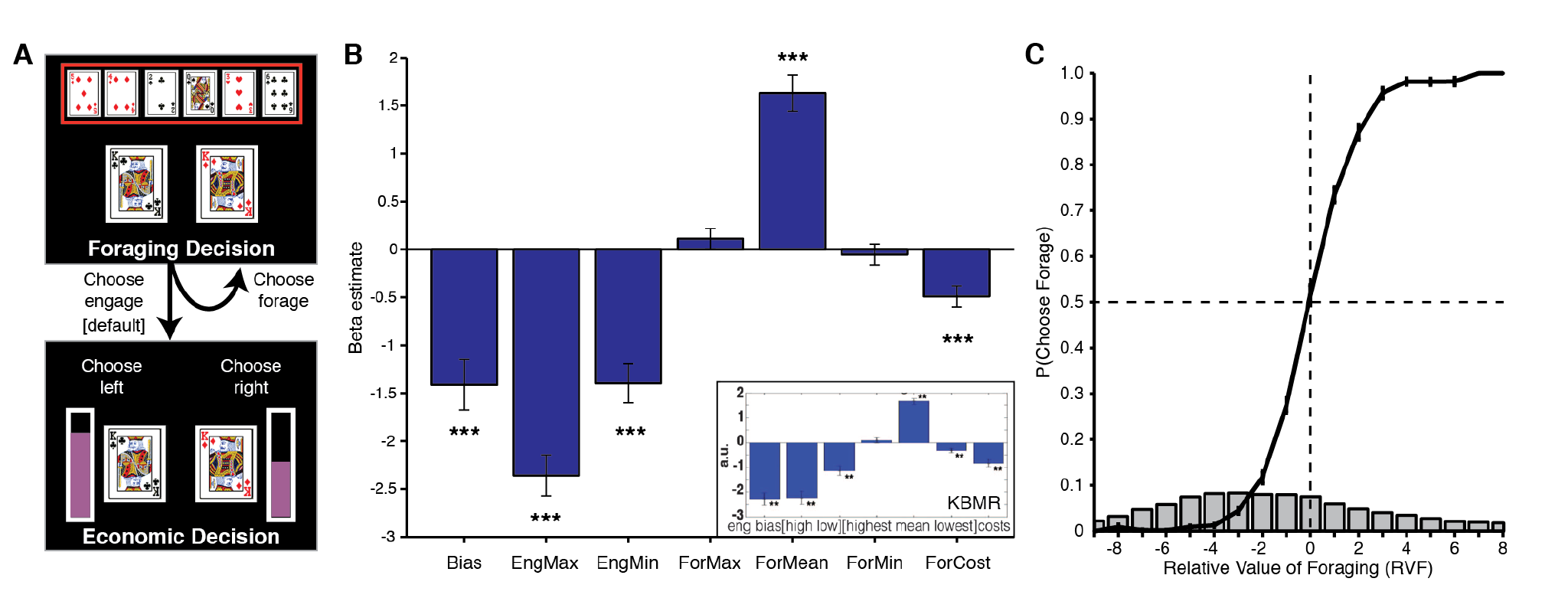
Behavior in foraging task. **A)** Participants made foraging-like decisions in Stage 1 of each trial, with playing cards indicating potential reward values, and the color of the search box indicating the cost of foraging. Once they decided to continue to Stage 2 (engage), they made an economic choice based on reward magnitudes and their probabilities (level of purple bar). **B)** Results of regressing forage choices in Stage 1 on each of the relevant decision variables (max/min of the engage pair, max/mean/min of the forage set, forage cost). Inset: Results from the equivalent regression in KBMR. **C)** Probability of choosing to forage (vs. engage) as the relative value of foraging (RVF) increased. RVF is calculated based on the subject-specific log-odds of choosing to forage (from regressions in Panel B). RVF therefore reflects a decision variable that has been fit to the choice data shown on the y-axis (rather than an independent canonical estimate). (See also Supplementary Figure 1A.) RVF values are binned (for display purposes only), and the gray bars represent a histogram of trial frequencies across RVF bins. This plot also excludes RVF bins with less than an average of five trials per subject. *** *p* < 0.005.

### Neuroimaging results: dACC activity continues to be accounted for by choice difficulty but not foraging value

Replicating KBMR and our previous study, we found a significant effect of RVF on dACC activity when focusing on a subset of trials that approximated those used in KBMR (i.e., trials whose RVF values predominantly favor engaging; Figure 2A) (GLM 1). However, this effect diminishes and then reverses as we test subsets that favor progressively higher values of RVF. Similarly, when including all trials, we find a significant positive effect of RVF for trials in which the participant chooses to engage but find a significant negative effect of RVF for trials in which the participant chooses to forage (Figure 2B; GLM 2). In other words, the effect of RVF seems to depend on whether RVF is approaching or diverging from indifference (i.e., whether the trial is increasing or decreasing in difficulty). Accordingly, when including RVF and choice difficulty (-|RVF|) in the same GLM, controlling for (log) RT (GLM 5), we find a significant effect of choice difficulty (*t*(30) = 5.8, *p*<0.001) but not RVF (*t*(30) = 0.34, *p*=0.73; Figure 2C). The effect of RVF also did not appear in the dACC ROI used in KBMR (x=4, y=28, z=30; *t*(30)<0.48, *p*>0.60), nor did it appear in either ROI when using alternate formulations of RVF (KBMR's *search evidence* [RVF_KBMR_]: *t*(30)<0.54 [GLM 7]; KBMR’s *search value* [modeled simultaneously with engage value]: *t*(30)<-1.3 [GLM 8]).

**Figure 2.**
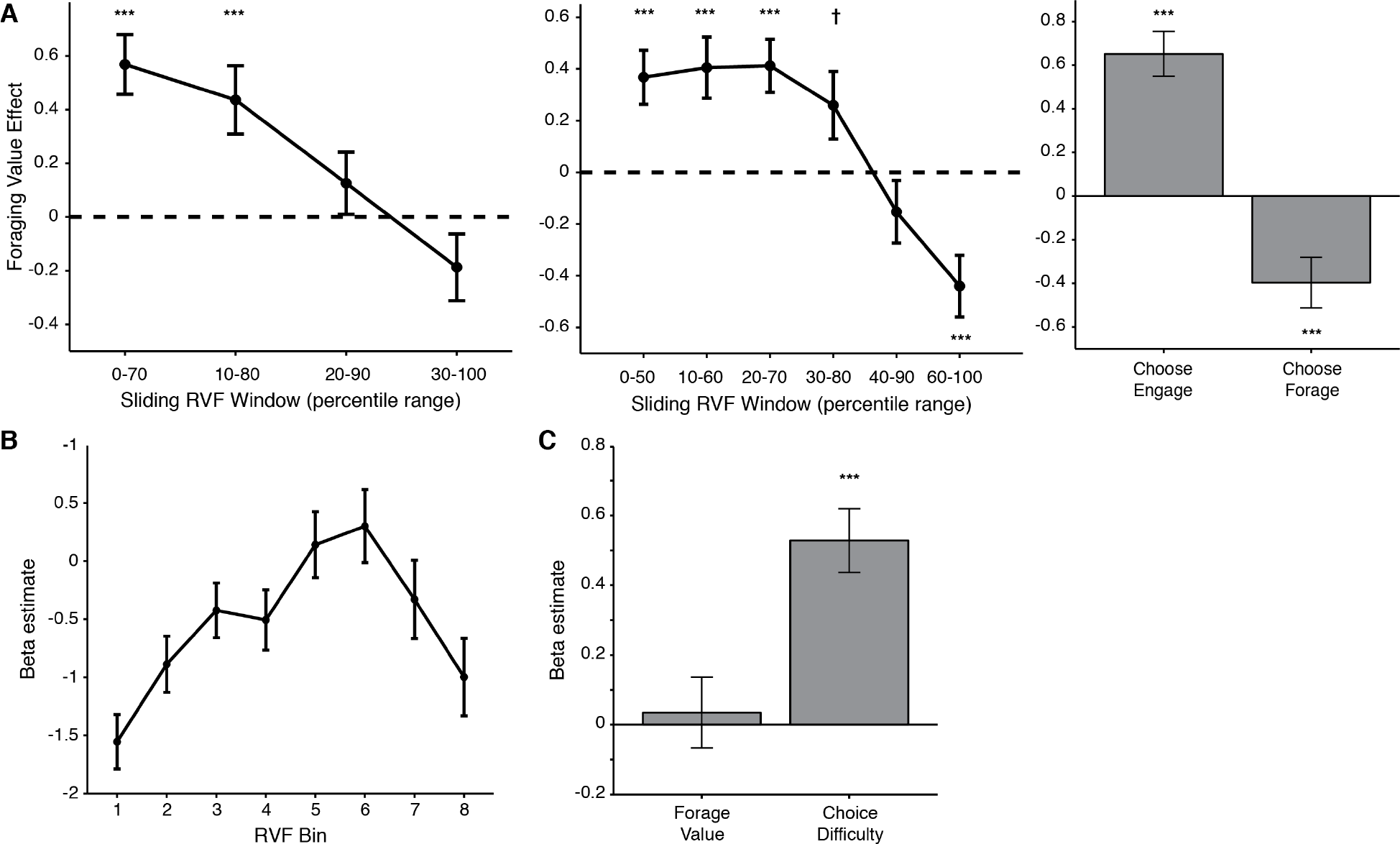
dACC ROI analyses replicate effects of foraging value only when confounded with difficulty. **A)** Left/Middle: Using an independent ROI defined in Shenhav et al. (2014), we tested for a parametric effect of increasing RVF within consecutive sliding windows that capture either 70% (left) or 50% (middle) of the foraging trials at a time. In both cases we replicate a significant positive effect of RVF on dACC activity for lower RVF values that decreases and eventually reverses at higher values. Right: Similarly, we see a significant positive (negative) effect when focusing on trials where the subject chose to engage (forage). **B)** A comparable analysis (GLM 3) shows that average activity in dACC is non-monotonically related to RVF, exhibiting an inverse U-shaped function. **C)** Accordingly, when including all forage trials in our analysis, we find no effect of foraging value (RVF) but instead a significant effect of choice difficulty (|RVF|). † *p* < 0.10, *** *p* < 0.005.

These effects are not specific to the dACC ROI just reported. Additional exploratory analyses failed to identify a region of dACC that tracks RVF across all trials, for instance any voxels that showed a positive relationship with RVF on both Choose Engage and Choose Forage trials (Figures 3–4, Supplementary Figure 6; GLMs 1–2, 5). Nor did dACC track RVF when we excluded the easiest choice trials (Supplementary Figure 3, GLM 5C; cf. Kolling et al., 2016). Instead we found that a wide swath of dorsal ACC was best accounted for by choice difficulty (e.g., the conjunction of a positive RVF effect on Choose Engage trials and a negative RVF
effect on Choose Forage trials).

**Figure 3.**
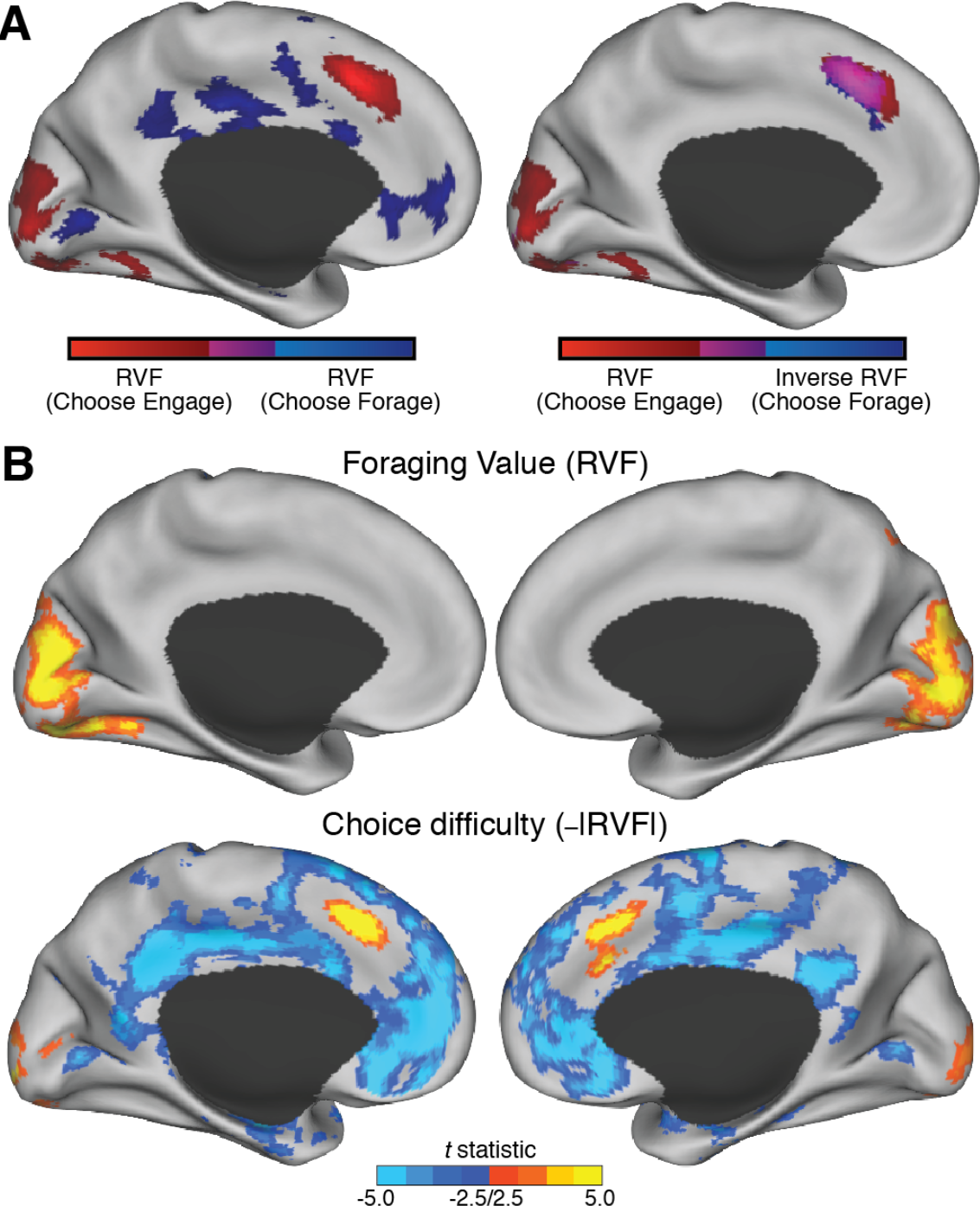
Whole-brain analyses only reveal dACC regions that track difficulty, not foraging value. **A)** Left: We tested for regions of dACC that showed a positive effect of RVF both when choosing to engage (red) and choosing to forage (blue). We did not find any overlap between these (magenta) in dACC. Right: However, we did find such overlap when looking for regions that positively tracked RVF on Choose Engage trials and negatively tracked RVF on Choose Forage trials. **B)** Across all trials, we fail to find any significant clusters in dACC that positively track RVF (top), but find that much of dACC is explained by difficulty (bottom). All statistical maps are shown at *p* < 0.01, and are extent-thresholded to achieve a FWE cluster-corrected *p* < 0.05. See also Supplementary Figures 1B and 2.

### Neuroimaging results: vmPFC activity is accounted for by relative chosen value in both task stages

The focus of the preceding analyses, and of our previous paper, is on the dACC and its potential role in foraging choices (Stage 1). However, an important finding from KBMR contrasted dACC’s putative role in foraging decisions with vmPFC's putative role in economic decisions (Stage 2). In particular, they reported that dACC activity better tracked the relative value of the foraging (non-default) option during Stage 1 than the relative value of the chosen option during Stage 2. They also reported that dACC activity was better tracked by choice difficulty (the relative value of the *unchosen* option) during Stage 2 than during Stage 1. We replicate these effects in our current data (Figure 5A; GLM 4). Our previous and current results suggest that dACC activity in both contrasts benefited from a narrow range of RVFs and a handicapped estimate of choice difficulty in the forage phase. However, our previous study did not address KBMR’s related finding, which suggested that activity in vmPFC was better tracked by the relative value of the chosen item during Stage 2 than during Stage 1 (Figure 5B); during Stage 1, vmPFC activity primarily tracked the value of the chosen option when it reflected the value of engaging (i.e., when participants chose to engage).

Based on these observations, KBMR argued that vmPFC activity is more directly linked to valuation and/or choice comparison during economic decisions (i.e., representing relative chosen value; Boorman, Behrens, Woolrich, & Rushworth, 2009; Boorman et al., 2013; Hunt et al., 2012), but during foraging decisions its role seems most closely tied to the relative value of *engaging*. They contrasted this with a dACC role in representing relative *unchosen* value (or difficulty) during economic decisions and the relative value of *foraging* in Stage 1. The striking similarities between the symmetric inversions in both stages, combined with our finding that the effect of foraging value in ACC masked a difficulty effect, led us to hypothesize that vmPFC activity in Stage 1 might have in fact reflected the opposite of choice difficulty (i.e., choice ease, or relative chosen value) rather than the opposite of forage value (i.e., engage value). Consistent with this, we found that activity in vmPFC tracked relative engage value (i.e., negatively tracked RVF) on a similar range of trials as in KBMR (when dACC foraging value effects were most positive), but that this effect reversed in concert with dACC, so that vmPFC *positively* tracked forage value at the uppermost window of our sliding foraging value analysis (Figure 4).

**Figure 4.**
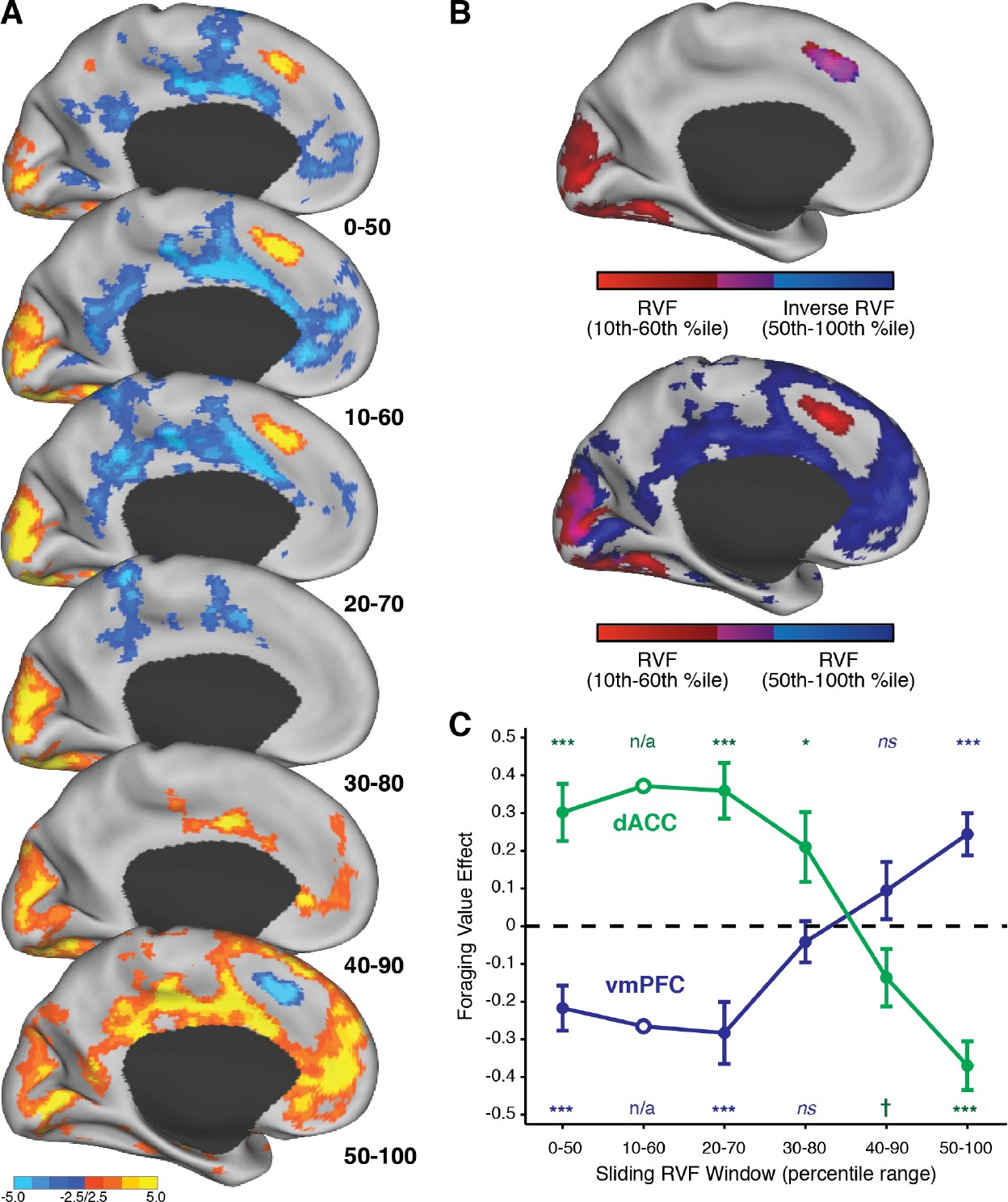
Inverse effects of foraging value in dACC versus vmPFC**. A)** Whole-brain map for each of the RVF windows shown at the bottom of Figure 2A. As these windows increase, RVF effects across dACC go from positive to negative. Over these same intervals, RVF effects in vmPFC go from negative to positive. See also Supplementary Figure 4A. **B)** Top:Regions of dACC are identified for the overlap of positive RVF effects in the 10^th^–60^th^ percentile window and negative RVF effects in the 50^th^–100^th^ percentile window. Bottom: No such overlap is seen in dACC when examining positive RVF effects in both windows, but instead the positive RVF effect in dACC appears to transfer to vmPFC. **C)** We tested whether the region of dACC (vmPFC) that show the strongest positive (negative) foraging value effect in low RVF foraging trials (open circles; SEM omitted due to circularity) would maintain such an encoding at higher RVFs. Instead we see that foraging value encoding reverses in both regions. See also Supplementary Figure 4B. † *p* < 0.10, * *p* < 0.05, ** *p* < 0.01, *** *p* < 0.005.

Collectively, our results in both dACC and vmPFC suggest unity rather than dissociation between the decision mechanisms for the foraging and engage stages of this task. Consistent with this, we found that choice difficulty accounted for activity in dACC to a similar degree and across a similar extent of cortex in both stages (Figure 5C, top; GLM 6). (Note that this was also true of other regions typically coactivated with dACC, including lateral parietal and prefrontal cortices.) The same was true of vmPFC (and networked regions, including posterior cingulate cortex) with respect to the ease of a given choice (Figure 5C, bottom).

**Figure 5.**
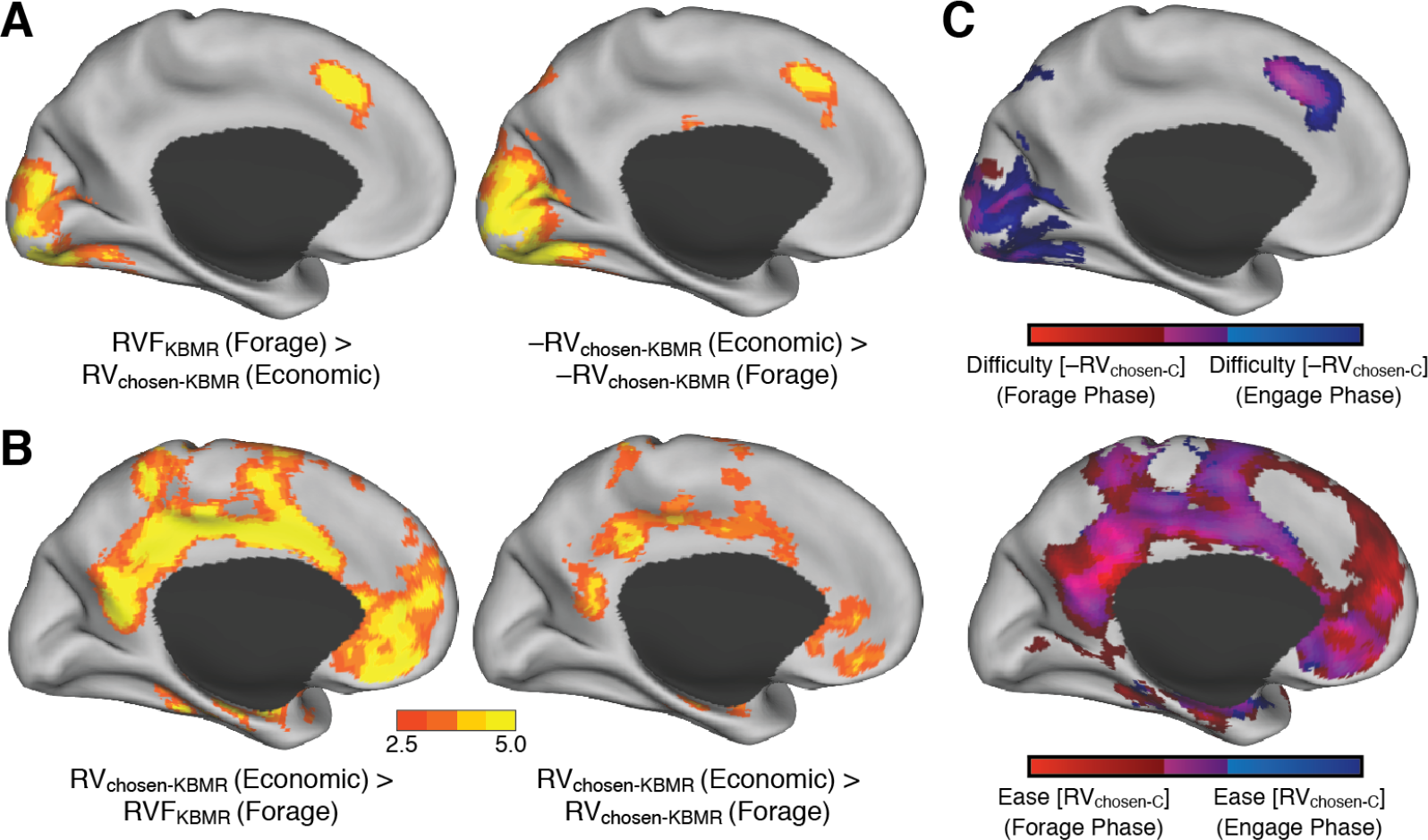
dACC and vmPFC perform similarly opposing roles for both foraging and economic choice. **A)** As in KBMR, when we test a narrow range of foraging values (0 –70^t^ percentile), using a canonical estimate of RVF uncorrected for bias (RVFkbmr), we find that dACC activity is better explained by foraging value in Stage 1 than by the relative value of the chosen option (RVchosen-KBMR) in Stage 2 (left), and it is better explained by relative value of the unchosen option (-RVchosen_-_KBMR) in Stage 2 than in Stage 1 (right). **B)** Conversely, we find that vmPFC is better explained by RVchosen_-KBMR_ in Stage 2 than either foraging value (left) or RVchosen_-KBMR_ (right) in Stage 1. To emulate analyses in KBMR, analyses shown in Panels A and B exclude four participants who did not choose the forage option at least eight times within this subset of trials. **C)** Using all trials and corrected estimates of RV, we find that activity in dACC (top) versus vmPFC (bottom) in both stages is well-explained by choice difficulty versus ease. Note that these maps represent correlates of ±RV_Chosen-C_, to parallel those shown in Panels A-B. However, qualitatively identical patterns are found for l|RVFC|, the measure of ease/difficulty used elsewhere in the main text, given that |RVF_-C_| and RV_Chosen-C_ are very highly correlated. See also Supplementary Figure 5.

A re-analysis of Experiment 2 from Shenhav et al. (2014) identified the same patterns in both regions, something we had only previously established within dACC. Specifically, using the same RVFc measure and dACC/vmPFC ROIs as in the current study, we find a significant effect of choice difficulty in dACC in both choice stages of this previous study (*t*_forage_(13) = 5.9, *t*_economic_(13) = 4.8, p<0.001; GLM 5B). We also find a significant effect of choice ease in vmPFC in both stages (*t*_forage_(13) = 3.1, *t*_economic_(13) = 3.6, *p*<0.01). These analyses control for RVF, which does not correlate with activity in either region in this earlier study (|*t*(13)| < 0.60, *p*>0.55). Notably, these vmPFC activations were weak and non-significant when testing for correlates of choice ease based on the RVF measure that was at the focus of our previous study (|*t*(13)| < 1.2, *p*>0.25). This RVF measure did correct for a given subject’s default foraging biases, but the value estimates being corrected were based on KBMR’s canonical formula for RVF rather than on within-subject choice regressions.

## Discussion

Our current findings replicate and extend our previous ones. We once again find that dACC activity is better explained by choice difficulty than foraging value, and we build on this result in two ways. First, with greater power (N=31 vs. N=14) and a task more closely matched to KBMR, we provide more conclusive evidence against the hypothesis that dACC tracks both choice difficulty and foraging value. Second, we shed new light on the role of vmPFC in this foraging task, showing that this region is engaged under similar conditions (increasingly easy choices) in both stages of the task. As was the case for dACC, the vmPFC result is obscured when a more limited range of foraging values is used, leading to apparent differences between this region’s involvement across choice types. Overall, we find that neural activity in foraging suggests it recruits areas and involves computations qualitatively equivalent to those involved in traditional economic choice.

Our previous findings ruled out the possibility that dACC tracks the value of foraging instead of choice difficulty (i.e., that the former accounted for the latter), but left open the possibility that it tracked both. However, the current findings, taken together with those earlier ones, call into question whether the dACC tracks the value of foraging at all. In both studies we find that the same regions of dACC that showed a positive relationship with foraging value for the range of RVFs in which the easier choice was to engage (i.e., ones that best approximated KBMR’s design), show a robust *negative* association for the range of RVFs in which the easier choice is to forage. We note that in this respect our current study is the third rather than the second to show such an effect. Kolling and colleagues (2014) examined dACC activity while subjects made risky choices in order to accumulate reward toward a target. Their subjects had a baseline bias against risk, but were driven to make risky choices based on the relevant trial values. The study found that dACC showed a positive relationship with the variable that exerted the strongest influence on choice (‘risk pressure’) at lower values (when subjects were choosing the default, safe option), but this relationship decreased and/or reversed at higher values (when subjects were choosing the risky option). They did not find such a difference when examining a second variable that impacted choice (‘relative value of the risky option’) but this could be because this latter variable exerted a much smaller influence on choice than risk pressure (there was more than a three-fold difference in their weights in a logistic regression on risky choice).

We designed the current study to address gaps that remained between the designs of KBMR’s original task and the second experiment of our previous study. The most significant of these were an enforced response delay (omitted in our previous study to capture and model the full extent of RTs), and our previous use of entirely numeric stimuli (in order to capture a wider range of rewards). The absence of a response delay in our original study may have clouded our ability to detect evaluative signals in dACC, including the possibility of ones reflecting the value of foraging as proposed by KBMR. Furthermore, the use of purely numeric stimuli in Experiment 2 of our previous study may have more readily enabled mathematical operations and, in doing so, could have decreased our sensitivity to evaluative signals (e.g. a foraging value signal over and above a difficulty-related signal). In addition to enforcing a response delay, the current task used playing card stimuli, which are ranked by both suit and number/face (including an out-of-order number, the ace). In addition to having been shown to have evaluative significance in previous neuroscience experiments (e.g., Preuschoff, Quartz, & Bossaerts, 2008; Rudorf, Preuschoff, & Weber, 2012), these cards were intended to render explicit mathematical operations difficult, particularly given that participants were simultaneously evaluating search costs that were conveyed by the color of the forage option frame. As in our previous study, these stimuli were chosen to enable sufficient variability in foraging rewards (52 cards), relative to the 12 unique stimulus-reward pairings that KBMR’s participants learned and later recalled during their task. However, the fact that the current stimuli were not identical to KBMR’s raises three concerns, which we address in turn.

First, as noted above for our previous study, it is possible that our stimuli enabled participants to perform explicit mathematical operations in lieu of a more value-based decision process. The findings of our follow-up experiment provide strong evidence against such an assertion. We instructed participants to perform the arithmetic operations needed to explicitly calculate foraging value, using a comparable if less complex stimulus display than the foraging task (Supplementary Figure 8). Their behavior bore little resemblance to behavior on the foraging task (e.g., RTs were an order of magnitude greater) and, in conjunction with subsequent self-reports, suggested that participants performing the foraging task would have been very reluctant to resort to math to guide their foraging choices. Second, it is possible that participants in our study engaged in other forms of strategies or heuristics in addition to or instead of valuation.

Such a concern is impossible to rule out for either our current study involving playing cards or for KBMR’s original study involving unique abstract stimuli. In fact, when using those same stimuli in Experiment 1 of our previous study, subjects frequently reported strategies involving subsets of those stimuli. The key question is whether participants resorted to such strategies more in the current study. The fact that our participants and KBMR’s exhibited highly similar patterns of behavior (Supplementary Figure 7) and foraging value-related BOLD activity (when constraining neural data to similar trial ranges; Figures 4–5) suggests that this is unlikely. Finally, it is possible that in order to evoke foraging value-specific signals in dACC one must use stimuli whose ordinal relationships are unknown prior to the experiment. We of course cannot rule this out as a reason we were unable to find such signals in the current study, but we note that revising a foraging account to make such a specific prediction would preclude the support of other findings that have been brought to bear on this account (Hayden et al., 2011; Kolling et al., 2014; Mobbs et al., 2013), and would arguably be in tension with descriptions of rewards available in real-world foraging environments (which typically have ordinal properties like size and color that indicate amount and quality of the reward to be gained).

While these findings collectively paint a picture inconsistent with a foraging account of dACC, it is important to note that the difficulty-related activations we observed lend themselves to a number of possible interpretations. These include accounts of dACC as monitoring for cognitive demands such as conflict/uncertainty (Botvinick, Braver, Barch, Carter, & Cohen, 2001; Cavanagh & Frank, 2014), error likelihood (Brown & Braver, 2005), and deviations from predicted response-outcome associations (Alexander & Brown, 2011); indicating the aversiveness of exerting the associated cognitive effort (Botvinick, 2007); explicitly comparing between candidate actions (e.g., Hare, Schultz, Camerer, O'Doherty, & Rangel, 2011); and regulating online control processes (Dosenbach et al., 2006; Posner, Petersen, Fox, & Raichle, 1988; Power & Petersen, 2013). In line with a number of these accounts, we recently proposed that dACC integrates control-relevant values (including factors such as reward, conflict, and error likelihood) in order to make adjustments to candidate control signals (Shenhav, Botvinick, & Cohen, 2013). In this setting, one of multiple potentially relevant control signals is the decision threshold for the current and future trials, adjustments of which have been found to be triggered by current trial conflict (i.e., difficulty) and mediated by dACC and surrounding regions (Cavanagh & Frank, 2014; Cavanagh et al., 2011; Danielmeier, Eichele, Forstmann, Tittgemeyer, & Ullsperger, 2011; Frank et al., 2015; Kerns et al., 2004). Our findings also do not rule out the possibility that dACC activity will in other instances track the likelihood of switching rather than sticking with one’s current strategy-as has been observed in numerous studies of default override (see Shenhav et al., 2013)-over and above signals related to choice difficulty. However, our results do suggest that a more parsimonious interpretation of such findings would first focus on the demands or aversiveness of exerting control to override a bias, rather than on the reward value of the state being switched to.

Another recent study has questioned the necessity of dACC for foraging valuation by showing that this region does not track an analogous value signal in a delay of gratification paradigm involving recurring stay/switch decisions (McGuire & Kable, 2015). Instead, this study found that vmPFC played the most prominent evaluative role for these decisions (tracking the value of persisting toward the delayed reward). The authors concluded that vmPFC may therefore mediate evaluations in both foraging and traditional economic choice. Our study tested this assumption directly within the same foraging task that was previously used to suggest otherwise. We confirmed that vmPFC tracked relative chosen value similarly in both task stages. As is the case for our (inverse) findings in dACC, there are a number of possible explanations for this correlate of vmPFC activity, over and above the salient possibility that these activations reflect the output of a choice comparison process (Boorman et al., 2009; Boorman et al., 2013; Hunt et al., 2012). First, it may be that vmPFC activity is in fact tracking ease of choice or some utility associated therewith (cf. Boorman et al., 2009), such as the reward value associated with increased cognitive fluency (Winkielman, Schwarz, Fazendeiro, & Reber, 2003). Similarly, it may be the case that this region is tracking confidence in one's decision (possibly in conjunction with value-based comparison), as has been reported previously (De Martino, Fleming, Garrett, & Dolan, 2013; Lebreton, Abitbol, Daunizeau, & Pessiglione, 2015). A final possibility is that this region is not tracking ease or confidence per se, but a subtle byproduct thereof: decreased time spent on task. Specifically, it is possible that a greater proportion of the imposed delay period in this study and in KBMR was filled with task-unrelated thought when participants engaged with an easier choice, leading to greater representation of regions of the so-called “task-negative” or default mode network (Buckner, Andrews-Hanna, & Schacter, 2008), including vmPFC. While this is difficult to rule out in the current study, our finding of similar patterns of vmPFC activity in our previous experiment (which omitted an imposed delay) counts against this hypothesis. These possibilities notwithstanding, our results at least affirm McGuire & Kable’s conclusion that the vmPFC’s role during foraging choices does not differ fundamentally from its role during traditional economic choices.

Despite the remaining ambiguities regarding their individual functions, the patterns in these two regions together invite the integrative interpretation that foraging-like and more traditional economic choice share more similarities than differences at the level of mechanism. Moving forward, we recommend that experimenters trying to identify and dissect these similarities (a) account for all decision-relevant factors when determining the relative ease or difficulty of a given decision and (b) sample the space of decision values as widely as possible. We have proposed a few ways for doing the former in our current and previous paper, in each case incorporating default choice biases (e.g., towards exploitation of a safe option) into our estimate of choice difficulty. However, each of these approaches to modeling choice difficulty makes assumptions that need to be verified as appropriate for the given choice setting. For instance, by employing a logistic regression we assume that the thresholds for the two responses are equidistant from the starting point of the decision process, an assumption that may be violated in decisions that require overriding a default (i.e., if the decision-maker sets a lower threshold for choosing the default rather than the non-default option). Additionally, we assumed that subjects evaluate all of their choice options simultaneously rather than, for instance, only evaluating the foraging options if the engage options fail to meet some initial threshold. One way to evaluate these assumptions is to examine RT patterns across conditions of interest, provided that these RTs reflect the complete decision process (rather than, for instance, being truncated by an imposed delay period). These notes of caution aside, our findings suggest that vmPFC and dACC functions are similar irrespective of whether an individual is engaged in a foraging or an economic choice.

## Author Note

The authors are grateful to Gus Baker for assistance in data collection. This work was funded by a postdoctoral fellowship from the CV Starr Foundation (A.S.) and by the John Templeton Foundation. The authors declare no competing financial interests.

## Supplementary Figures

**Supplementary Figure 1.**
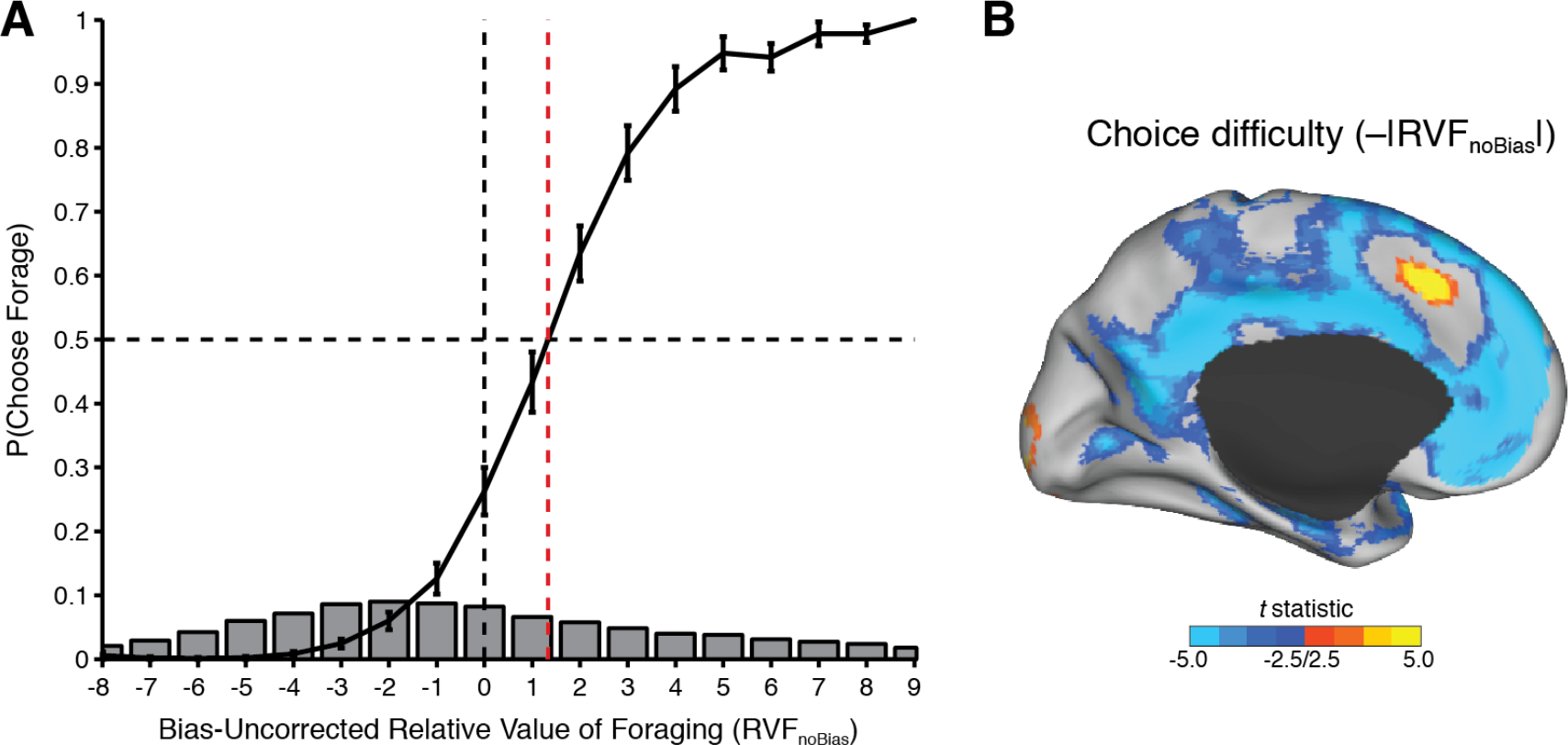
**Distribution of choices relative to an uncorrected estimate of RVF**. We generated a trial-by-trial version of RVF that includes all components of the logistic regression (as in Fig. 1C) except for the intercept term (RVF_noBias_). **A)** Whereas choice indifference (50%) occurs at RVF = 0 in Fig. 1C, this plot shows that the indifference point occurs to the right of RVF_noBias_ = 0, as expected. This plot excludes (bias-uncorrected) RVF bins with less than an average of five trials per subject. **B)** Because we designed the experiment to to sample broadly around either indifference point, choice difficulty based on this alternate version of RVF continues to correlate with BOLD activity in dACC and vmPFC (and associated networks).

**Supplementary Figure 2.**
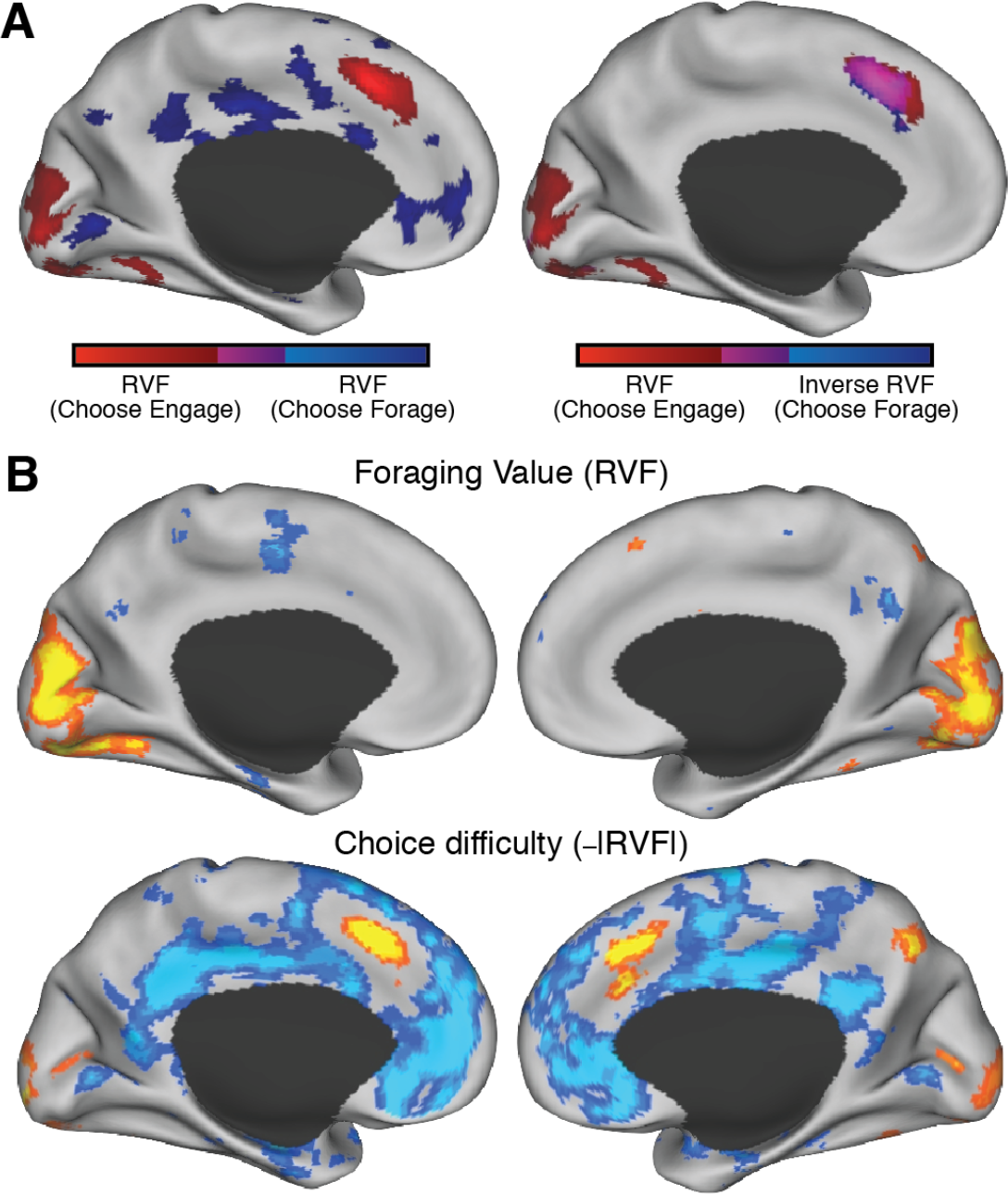
Panels are identical to Figure 3, but maps are shown at a more
liberal threshold (voxelwise p < 0.01, uncorrected for multiple comparisons).

**Supplementary Figure 3.**
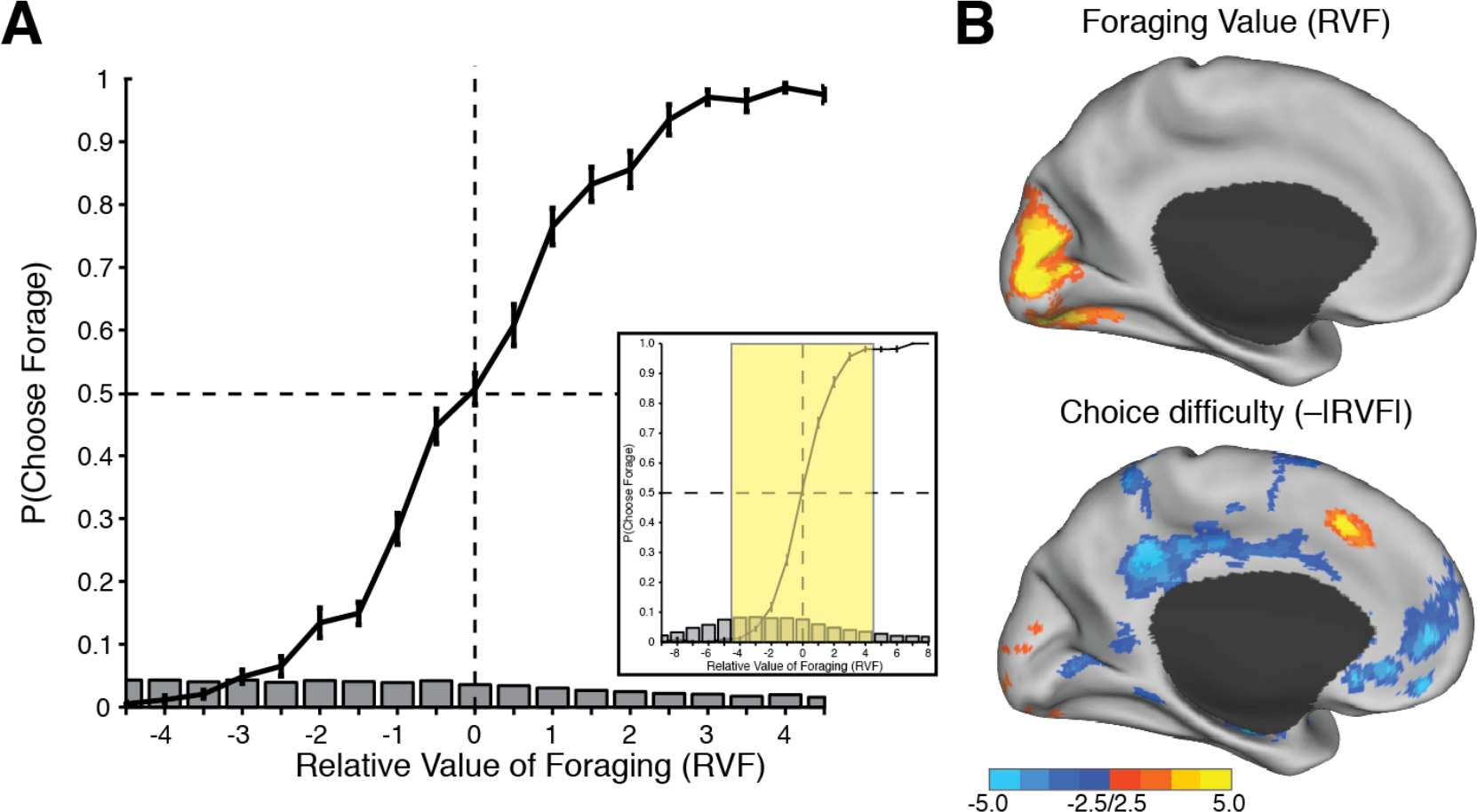
To test whether our findings change when excluding especially easy choices, we repeated our key analysis (GLM 5) after excluding trials with |RVF| (i.e., log-odds) values greater than 4.5 (GLM 5C). **A)** Choice curve for this narrower set of trials. Inset: choice curve for all trials (Fig. 1C), with the relevant window highlighted. **B)** Whole-brain corrected statistical maps show that we continue to find choice difficulty-related effects in dACC and vmPFC, but RVF effects still do not emerge in dACC (compare Fig. 3B).

**Supplementary Figure 4.**
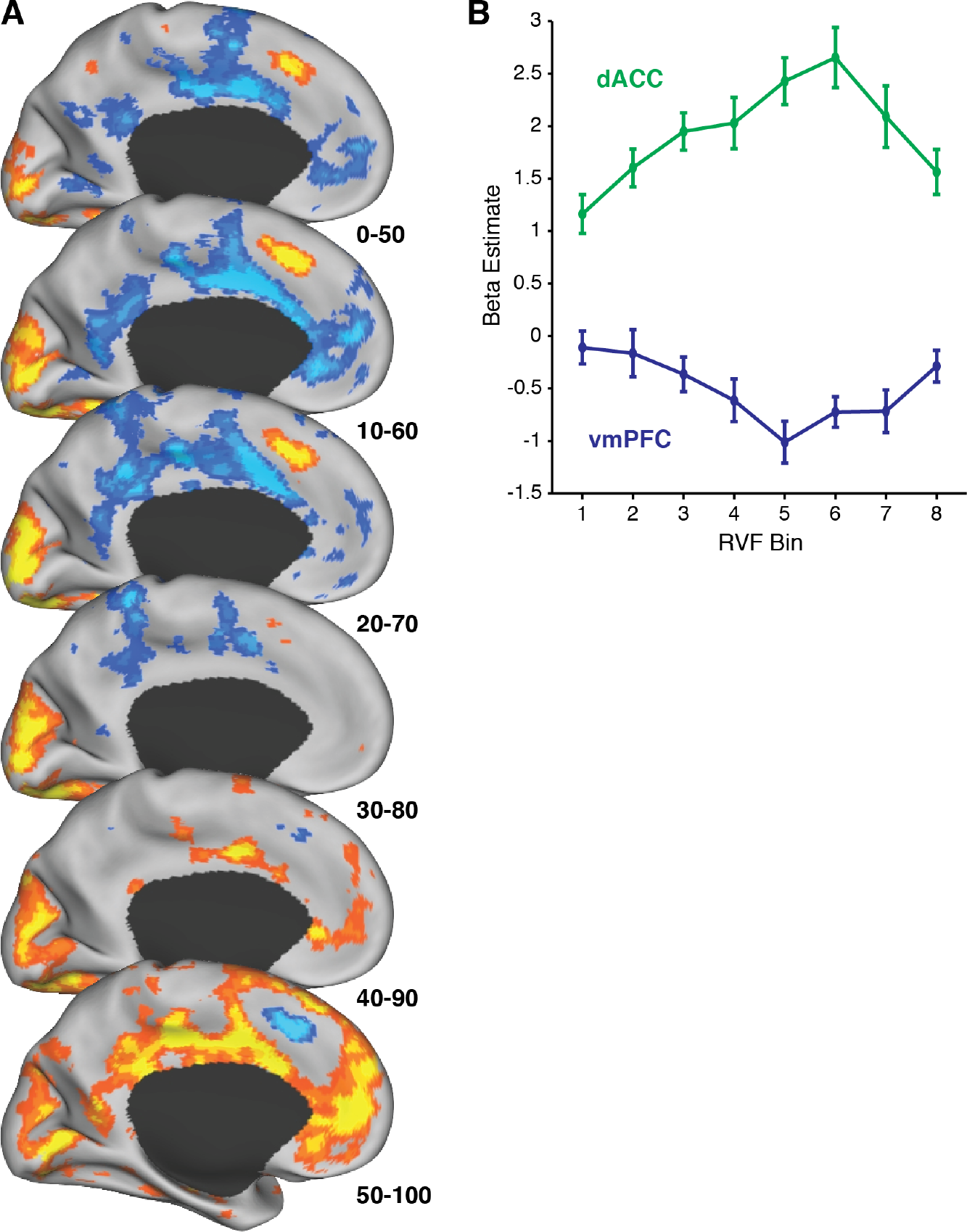
A) Identical to Fig. 4A, but maps are shown at a more liberal threshold (voxelwise *p* < 0.01, uncorrected for multiple comparisons). **B)** Average BOLD activity by RVF bin for ROIs shown in Fig. 4C (same analysis procedure as in Fig. 2B). Note that these ROIs were intentionally defined in a non-independent manner (to bias the detection of the initially positive [dACC] or negative [vmPFC] slope), and should therefore be interpreted with caution.

**Supplementary Figure 5.**
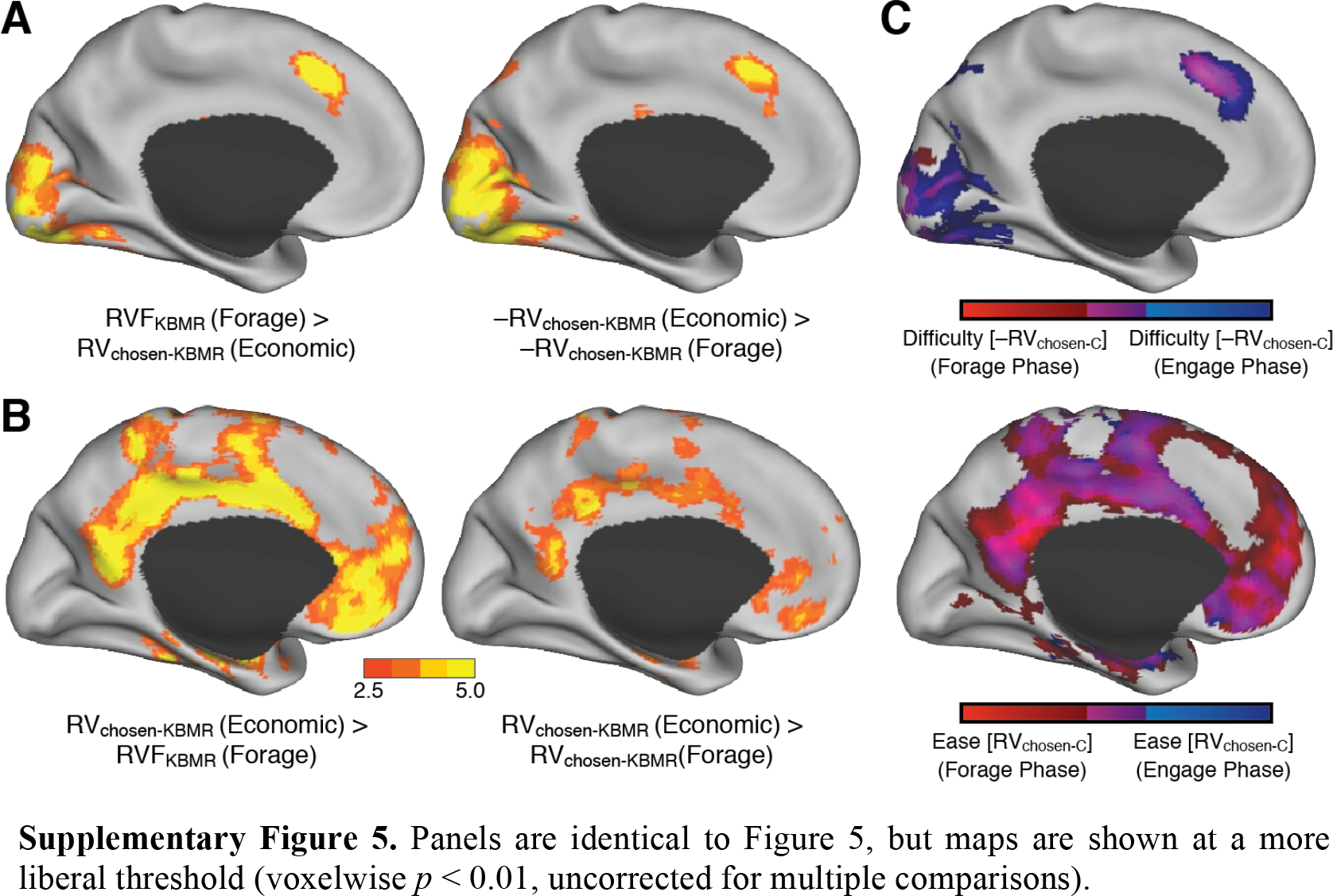
Panels are identical to Figure 5, but maps are shown at a more liberal threshold (voxelwise *p* < 0.01, uncorrected for multiple comparisons).

**Supplementary Figure 6.**
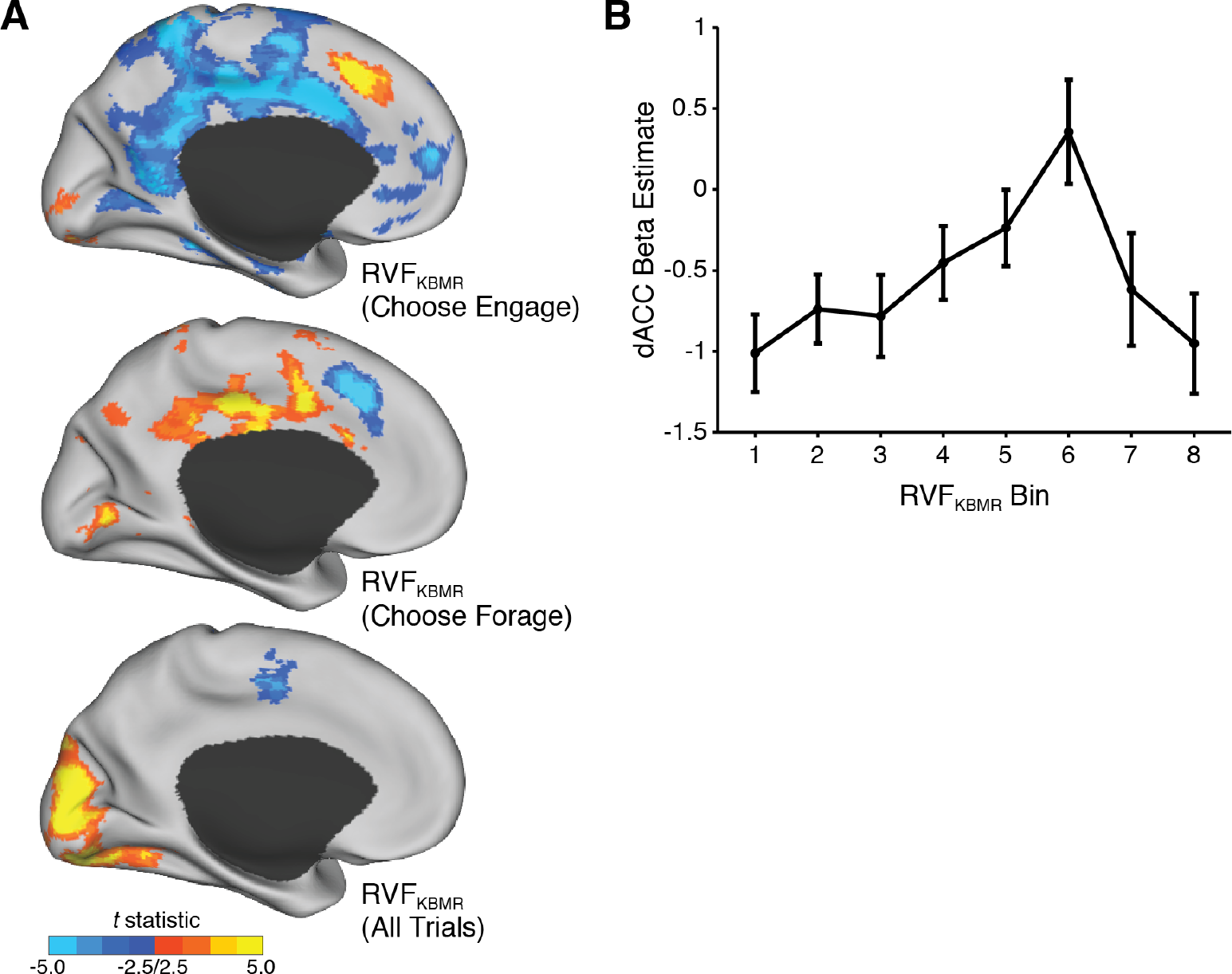
**A)** Top/middle: Parametric correlates of RVF_KBMR_ when subjects chose the default (top) and when they chose the non-default (middle) (GLM 2B; compare Fig. 3A). Bottom: No regions in dACC correlate with RVF_KBMR_ across all trials (GLM 7; compare Fig. 3B, top). All maps are whole-brain cluster-corrected. **B)** Average BOLD activity in dACC ROI (same ROI as in Fig. 2) by RVFKBMR bin.

**Supplementary Figure 7.**
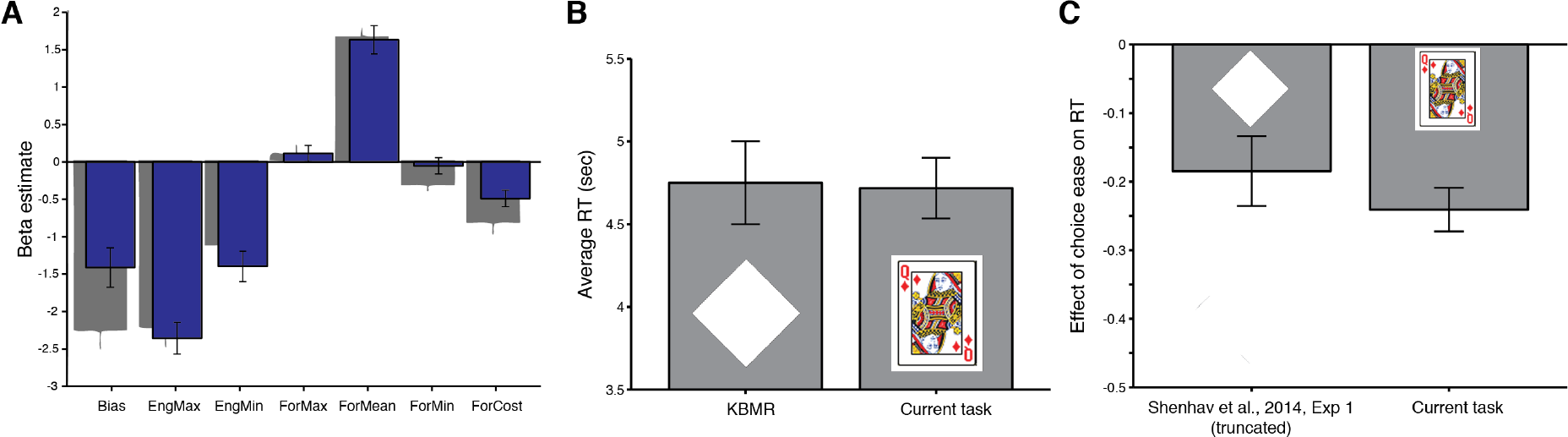
**Comparison of behavior using current stimuli versus novel stimuli used in KBRM and in Experiment 1 of Shenhav et al. (2014)**. **A)** Participants in the current study (blue) weighted foraging choice components similarly to participants in KBMR (gray). Gray bars adapted from Kolling et al. (2012). **B)** Average RTs in the current study (4.72s, SEM: 0.18) did not differ significantly from KBMR (4.75s, SEM: 0.25) (*t*(47) = 0.08, *p* = 0.93). Representative stimuli are shown for each study. **C)** The speeding effect of choice ease (absolute log-odds) on RT was not weaker in the current study (regression coeff = –0.24, SEM: 0.03) than in Shenhav et al., (2014, Expt 1; coeff = –0.18, SEM: 0.05); only the opposite trend is observed: *t*(44) = –1.08, *p* = 0.28. Because this previous study used a free-response paradigm, we have truncated RTs from that study (set the minimum to 2.7s) to approximate the RT distribution from the current study. While this provides a more appropriate comparison, we note that the choice ease effect for the untruncated RT distribution (–0.27, SEM: 0.05) is also not significantly larger than the current study (–0.24, SEM: 0.03) (*t*(44) = 0.54, *p* = 0.59). Caution is of course warranted when comparing across studies that differ in task parameters, but these findings collectively argue against the possibility that the stimuli in the current study elicited qualitatively different behavior than those used in KBMR.

**Supplementary Figure 8.**
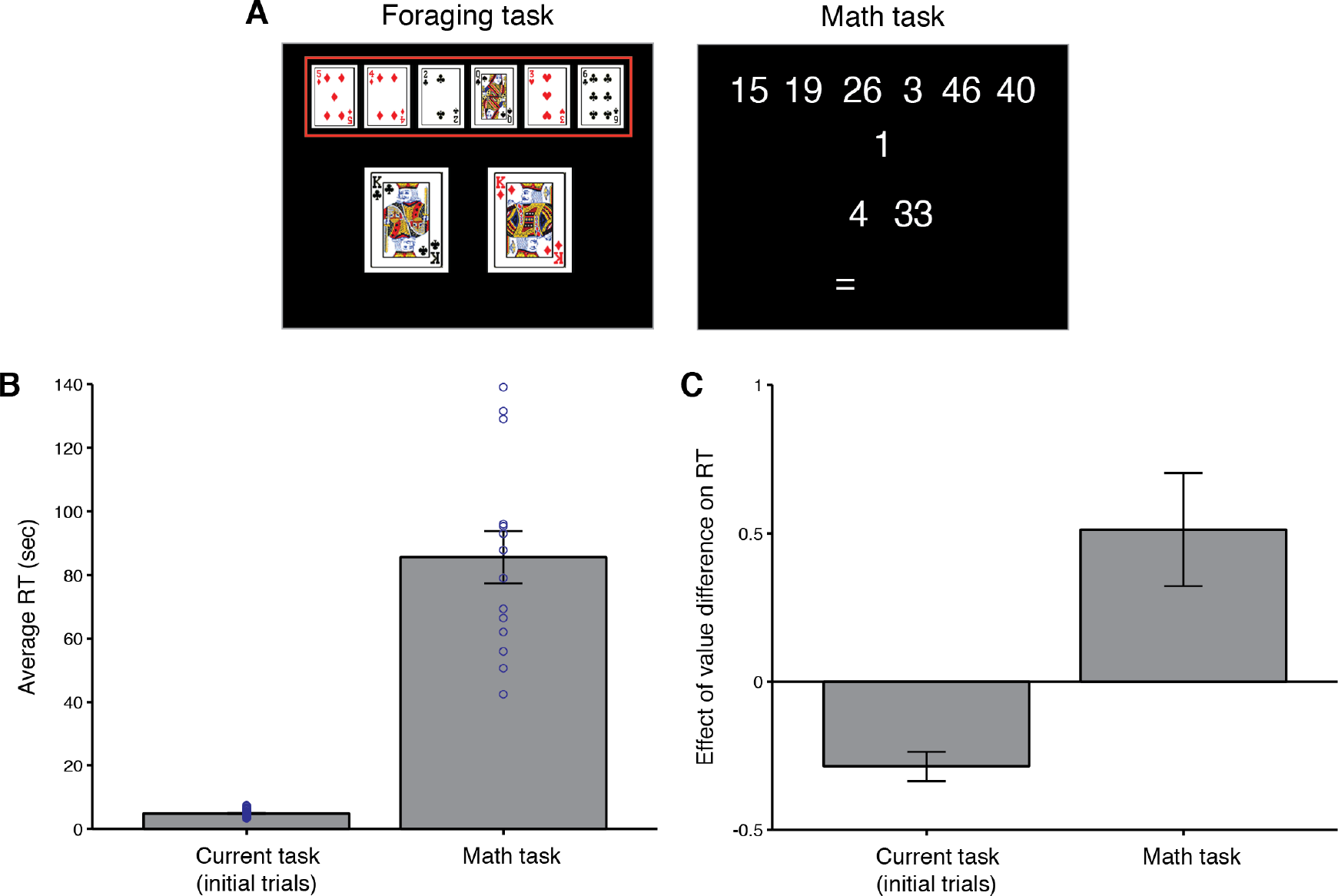
A) Participants in an independent sample (N=14) viewed a numbers-only equivalent of the foraging display and were instructed to explicitly calculate the equivalent of relative foraging value (i.e., the difference between the average of the top six numbers and the average of the bottom two numbers, minus the middle number). Participants typed their answer, and the response time was recorded (conservatively) as the time that they started typing (i.e., pressed the first key in their response). Participants performed the task until they either completed 60 trials or 45 minutes had passed, whichever came first. **B)** Average RTs in the math task (85.6s, SEM: 8.2) were substantially slower than in the foraging task used in the current study (4.81s, SEM: 0.19) (*t*(43) = 14.9, *p* < 0.0001). Scatter plots on both plots show individual participant average RTs, revealing that the slowest participant on the foraging task (M=7.4s) and the fastest participant on the math task (M=42.5s) differed by over 35 seconds. To provide a fairer comparison between tasks, we restrict all analyses here and in Panel C to the first 30 trials of the foraging task (the average number of trials completed by participants performing the math task. **C)** Whereas RTs decreased with value difference (absolute log-odds) in the foraging task (coeff = –0.29, SEM: 0.05, *p* < 0.001)-as in KBMR (Supplementary Figure 7)-they instead increased with value difference (absolute value of the correct response) in the math task (coeff = 0.51, SEM: 0.19, *p* < 0.05).

